# RNA degradation heavily impacts mRNA co-expression

**DOI:** 10.1101/2022.09.21.508820

**Authors:** Óscar García Blay, Pieter Verhagen, Benjamin Martin, Maike M.K. Hansen

## Abstract

Co-expression of genes measured with single-cell RNA sequencing is extensively utilized to understand the principles of gene regulation within and across cell types and species. It is assumed that the presence of correlation in gene expression values at the single-cell level demonstrates the existence of common regulatory mechanisms. However, the regulatory mechanisms that should lead to observed co-expression at an mRNA level often remain unexplored. Here we investigate the relationship between processes upstream and downstream of transcription (i.e., promoter architecture and coordination, DNA contact frequencies and mRNA degradation) and pairwise gene expression correlations at an mRNA level. We identify that differences in mRNA degradation (i.e., half-life) is a pivotal source of single-cell correlations in mRNA levels independently of the presence of common regulatory mechanisms. These findings reinforce the necessity of including post-transcriptional regulation mechanisms in the analysis of gene expression in mammalian cells.

## INTRODUCTION

The emergence of single cell analyses has unlocked an exciting new chapter for understanding gene regulation in cells. Specifically, single-cell RNA sequencing (scRNA-seq) is a powerful and widely implemented tool that allows for transcriptome-wide quantification of mRNA in thousands of individual cells (Hwang et al., 2018). One feature that can be extracted from scRNA-seq datasets is gene-to-gene co-expression (Eisen et al., 1998), which has been extensively used over the last years in a range of applications to gain quantitative insights into gene regulation of eukaryotic cells. For example, co-expression of genes has been described as a robust tool to identify cell types or states from scRNA-seq datasets (Crow and Gillis, 2018). Specifically, co-expression has been implemented in order to study cellular heterogeneity of tissues and organs in physiological (Aizarani et al., 2019; Andrews et al., 2022; Muraro et al., 2016; Payen et al., 2021; Travaglini et al., 2020) and pathological contexts (Esmaili et al., 2021; Han et al., 2021), as well as dynamic processes such as cell signaling or changes in cell identity during development (Foreman and Wollman, 2020; Qadir et al., 2020; Salehi et al., 2021). On the other hand, gene co-expression in scRNA-seq datasets has been exploited to extrapolate the functional principles of gene expression regulation (Desai et al., 2021) from cellular populations in the form of gene-regulatory networks (Matsumoto et al., 2017). The most common approach is the construction and analysis of co-expression networks from pairwise correlation measurements (Vivian Li and Li, 2021; Wang et al., 2021). Single-cell co-expression network analysis has thus led to the characterization of novel regulatory pathways (Xie et al., 2021), which in turn has improved our understanding of common gene regulation principles between cell types and species (Crow et al., 2022; Harris et al., 2021). Given the expansive application of single-cell co-expression analysis in both current and likely future studies, it is essential to understand the causality and limitations of observed correlations in scRNA-seq datasets.

In order to exploit co-expression networks to study gene regulation, a common assumption is that gene-to-gene correlation or anticorrelation indicates an underlying functional relationship (Eisen et al., 1998; Oliver, 2000). Traditionally, it has been assumed that such co-expression patterns arise from common regulatory mechanisms that are shared between respective genes. Since gene expression is a multi-step process (e.g., promoter toggling, transcription, RNA processing) (Ronen et al., 2002), there are different scenarios from which functional (anti)correlation between genes may emerge (Munsky et al., 2012). For instance, gene-to-gene (anti)correlation could either be caused by shared transcription factors commonly affecting the transcription of a group of genes, or due to post-transcriptional processes occurring downstream of transcription (Figure S1A). However, in higher eukaryotes, the balance between transcriptional and post-transcriptional events underlying mRNA co-expression remain elusive. Early bulk co-expression experiments have shown that promoter sharing can be a major source of gene co-expression (Gu et al., 2011). Yet, more recent single cell studies have shown that shared target-regulator relationships are unlikely to result in co-expression and that most--but definitely not all--shared transcription factors fail to enforce co-expressed behavior among target genes (Ribeiro et al., 2021; Yin et al., 2021). Solving the interplay between regulatory architectures and effective co-expression is key for novel strategies that apply combined approaches to study transcriptional regulation (Jeong et al., 2021).

Here, we sought to identify the parameters that contribute to correlation and anticorrelation in scRNA-seq data without enforcing previous regulatory structures. To this end, we integrated scRNA-seq data from mouse embryonic stem cells (mESCs) with other existing datasets from mESCs, including gene-to-gene contact frequencies in Hi-C data (Nora et al., 2017), promoter activity in intron seq-FISH data (Shah et al., 2018), and transcript-specific half-life measurements (Sharova et al., 2009). The analyses show that coordination in promoter activity or gene-to-gene contacts do not appear to be the major sources of gene-to-gene correlation in RNA expression levels. Remarkably, mRNA degradation emerges as clear contributor to the (anti)correlations (i.e., co-expression) measured with scRNA-seq.

## RESULTS

### Promoter coordination and gene-to-gene contacts do not explain mRNA co-expression

In order to identify possible causes for the (anti)correlation in mRNA levels we integrated several datasets. To this end, we combined single-cell mRNA abundance data (STAR METHODS) with 3 additional published datasets providing promoter activity information (Shah et al., 2018), gene-to-gene contact frequencies (Nora et al., 2017), and mRNA half-lives (Sharova et al., 2009) (Figure 1A and 1B), all 4 datasets from mESCs. In order to accurately compare all data modalities, we limited the number of genes analyzed to those included in all datasets, totaling 5277 genes. To extract (anti)correlation regimes we first performed gene-by-gene pairwise Pearson correlation analysis from scaled scRNA-seq read counts, followed by hierarchical clustering (Figure 1A). Pearson correlation is preferred over Spearman because highly expressed genes in a low number of cells display more accurate Pearson correlation values than Spearman correlation values (Figure S1B and S1C), as described in other studies (Vandenbon, 2022). To incorporate the promoter behavior of each gene into our analysis, we assumed two-state promoter toggling as an underlying mechanistic model for the analyzed promoters (Esmaili et al., 2021; Kepler and Elston, 2001; Raj et al., 2006; Weinberger et al., 2005). Eukaryotic promoters have been described to switch between at least two (Harper et al., 2011; Zenklusen et al., 2008) distinct states: one state that allows for mRNA production (referred to as the ON state) and another state that is not permissive for mRNA production (referred to as the OFF state). This toggling results in the discontinuous or bursty behavior that is associated with transcription in eukaryotes (Chong et al., 2014; Coulon et al., 2013; Golding et al., 2005; Suter et al., 2011). The effective ON and OFF states of each promoter can be defined by the presence or absence of intronic signal respectively (Bahar Halpern et al., 2015; Shah et al., 2018). With this information, we computed the likelihood that pairwise promoters are more coordinated (i.e., ON or OFF at the same time) or anti-coordinated (i.e., one promoter being ON while the other is OFF and *vice versa*) compared to what is expected from independent random promoter behavior (Figure 1A and 1B, green and purple respectively).

**Figure 1:**
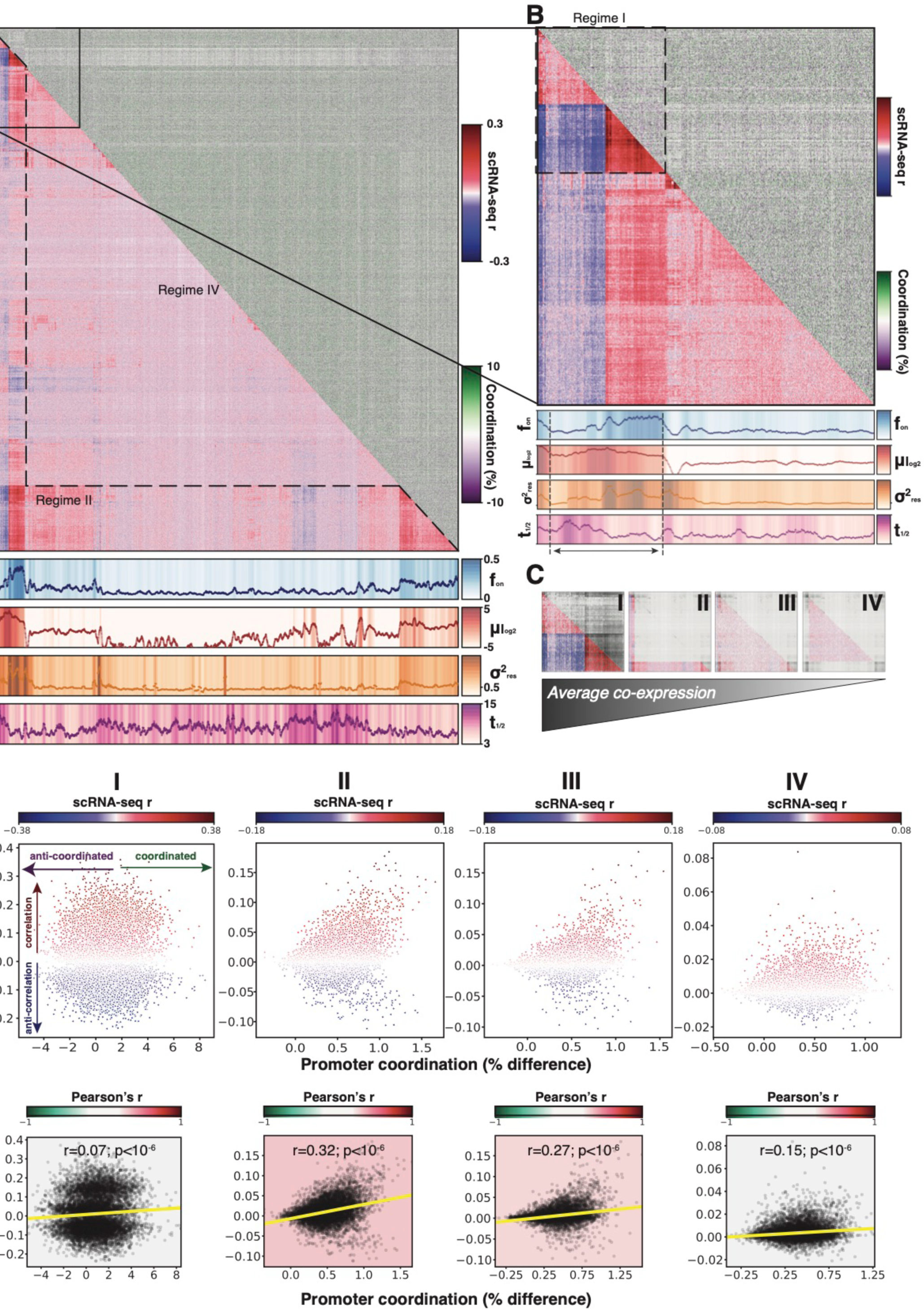
mRNA co-expression is not explained by promoter behavior. **(A-B)** Top: Combined matrix showing clustered pairwise scRNA-seq Pearson’s r coefficients (bottom half) and corresponding promoter (anti-)coordination scores (top). Red is positive Pearson’s r and blue is negative Pearson’s r. Green is more coordination and purple is more anti-coordination of promoters compared to what is expected from random promoter behavior. Bottom: Rolling averages for ON-ratio (*f*_*on*_), relative mean mRNA expression (μ_log2_), noise (∼Fano factor, σ ^2^_res_), and mRNA half-life (t_1/2_, hours) ranked in the same order as the clustered matrix, with the background color representing the respective values. **(C)** Four sampling regimes (I-IV) corresponding to decreasing average co-expression (i.e., scRNA-seq Pearson’s r) from left to right. **(D-E)** mRNA co-expression (i.e., scRNAseq Pearson’s r coefficient) versus average (anti-)coordination of pairwise promoters for regimes I, II, III and IV. **(D)** Color (blue to red) represent the average scRNAseq Pearson r coefficient. **(E)** Correlation between mRNA co-expression and promoter coordination, with linear regression line plotted in yellow and the background color corresponding to Pearson’s r correlation coefficient as a proxy for relationship strength.

To determine if mRNA correlation behavior (i.e., mRNA co-expression) originates at the promoter, we integrated the pairwise matrix of promoter coordination with the Pearson correlation matrix from the scRNA-seq dataset (Figure 1A and 1B, top). Surprisingly, the integrated matrices did not show any clear structure in the (anti-)coordination matrix when ranked according to the scRNA-seq r coefficient clustering, likely because promoter (anti-)coordination only deviated minimally from independent promoter behavior (Figure S1D). In order to gain deeper insights into the relationship of these two features that could be hidden in the visual representation of the data, we performed a quantitative analysis of 4 distinct correlation regimes with decreasing average co-expression values (Figure 1C): i) a small 387 by 387 gene cluster showing the highest and lowest values for positive and negative correlation respectively (Figure 1B and 1C, regime I); ii) an expanded regime including regime I in addition to a distant cluster with relatively high correlation values and all pairwise comparisons between these two clusters (Figure 1A and 1C, regime II); ii) the full 5277 by 5277 gene dataset (Figure 1A and C, regime III); iv) the central 4224 by 4226 gene regime where Pearson coefficients were close to 0 (Figure 1A, regime IV). Per regime (I-IV) we performed sub-sampling (see STAR Methods for more detail) and plotted the relationship between mRNA co-expression (i.e., scRNA-seq r) and promoter coordination (Figure 1D) and quantified the relationship by calculating Pearson’s correlation r as a proxy for the relationship strength (Figure 1D and 1E). The analysis showed a subtle positive association between promoter coordination and mRNA co-expression in regimes II and III (Figure 1E, red). Together, these data indicate that promoter coordination or anti-coordination does not generate a respective positive or negative correlation at an mRNA level.

To identify any hidden coregulation at a DNA level that was not captured by our promoter coordination analysis, we sought to discern if DNA contact frequencies were higher in gene clusters that are more (anti)correlated at an mRNA level (de Wit, 2020; Soler-Oliva et al., 2017). To this end, the same integration process (Figure S1E) and binned correlation analysis (Figure S1F and S1G) as for promoter coordination (Figure 1A-B and 1C-E respectively) was performed with Hi-C based DNA contact frequencies from a previously published dataset (Nora et al., 2017). Hi-C contact frequencies did not show any clear structure when plotted with respect to RNA co-expression (Figure S1E), nor did detailed quantitative analysis reveal a clear correlation between the Hi-C contact frequencies and mRNA co-expression (Figure S1F and S1G).

### mRNA half-life differences contribute to mRNA co-expression

Since neither promoter coordination nor DNA contact frequencies underlie mRNA co-expression behavior (Figure 1 and Figure S1), we considered whether differences in gene-specific kinetic parameters could be causing mRNA co-expression. From the intron seq-FISH dataset (Shah et al., 2018), we obtained the fraction of cells that are present in the ON state (i.e., *f*_*on*_) for each gene, which, assuming ergodicity coincides with the fraction of time a promoter is active (Dattani and Barahona, 2017; Desai et al., 2021). Numerically, *f*_*on*_ is a function of toggling rates--i.e., *f*_*on*_ = *K*_*ON*_*/[K*_*ON*_*+K*_*OFF*_]. On average, each promoter is on 12.5% of the time (i.e., *f*_*on*_ = 0.125), and each cell is expressing approximately 8% of the ∼10,000 genes analyzed at a given moment in time as previously described (Shah et al., 2018). The data (Shah et al., 2018) represents a population of cells where every cell has approximately the same number of genes active at a given timepoint, and the average promoter is in a bursty toggling regime (Munsky et al., 2012), where *K*_*ON*_*<K*_*OFF*_ (on average *K*_*OFF*_*=∼7xK*_*ON*_). These promoter toggling frequencies are in line with previous observations (Bahar Halpern et al., 2015; Hansen et al., 2018). To expand our analysis to post-transcriptional kinetic parameters, we included the half-life of each transcript (*t*_*1/2*_) from a third published dataset (Sharova et al., 2009).

First, in order to verify that these three separate datasets (mRNA co-expression, *f*_*on*_, and *t*_*1/2*_) could indeed be accurately compared, we determined if the relationship between mRNA expression, promoter toggling, and mRNA degradation behaved as expected. Overall, the fraction of time the promoter is in the ON state (*f*_*on*_) followed a similar trend as the mean mRNA abundance. This is especially evident when considering the trend across all ∼5000 genes (Figure 1A, blue and red respectively). This is expected, since the mean mRNA abundance (*μ*) is proportional to the time a promoter spends in the ON state (*μ ∝ f*_*on*_.*k*_*tx*_; where *k*_*tx*_ is the transcription rate) (Munsky et al., 2012). We next quantified the strength of the association between mean mRNA abundance and the fraction of time a promoter spends in the ON state (Figure 2A) or mRNA half-life (Figure 2B) as a function of mean expression. At low mRNA expression there is a strong positive association between mean mRNA abundance and the fraction of time a promoter spends in the ON state (*f*_*on*_). This association exhibits an overall decrease with increased mRNA expression levels. In other words, for genes with higher mRNA abundance, mean mRNA expression displays a much weaker association with promoter toggling frequencies than for genes with low mRNA abundance, indicating that other processes play a more dominant role. Interestingly, genes that show a decreased association between the fraction of time a promoter spends in the ON state (*f*_*on*_) and mean mRNA abundance, show a corresponding increase in association of mRNA half-life (*t*_*1/2*_) with mean mRNA abundance (Figure 2A and 2B, grey shaded area). This is in accordance with what is expected from the relationship of mean mRNA abundance, promoter toggling and mRNA degradation (Figure S2A) (Hansen et al., 2018; Munsky et al., 2012). Furthermore, the time a promoter spends in the ON state (*f*_*on*_) follows a similar trend as noise (*α*^*2*^_*res*_, equivalent to the Fano factor, see STAR Methods for more details), which is most apparent when considering the inset of ∼900 genes (Figure 1B, blue and orange respectively). We therefore quantified the strength of association between mRNA noise and the fraction of time a promoter spends in the ON state (Figure S2B). As expected from known dependence of noise on promoter toggling rates (Dar et al., 2012), mRNA noise is strongly associated with the fraction of time the respective promoter spends in the ON state (Figure S2A and S2B) with this association becoming stronger at higher mean mRNA abundances. Taken together, these results demonstrate that data obtained from different studies can reliably be compared, and relationships quantified.

**Figure 2:**
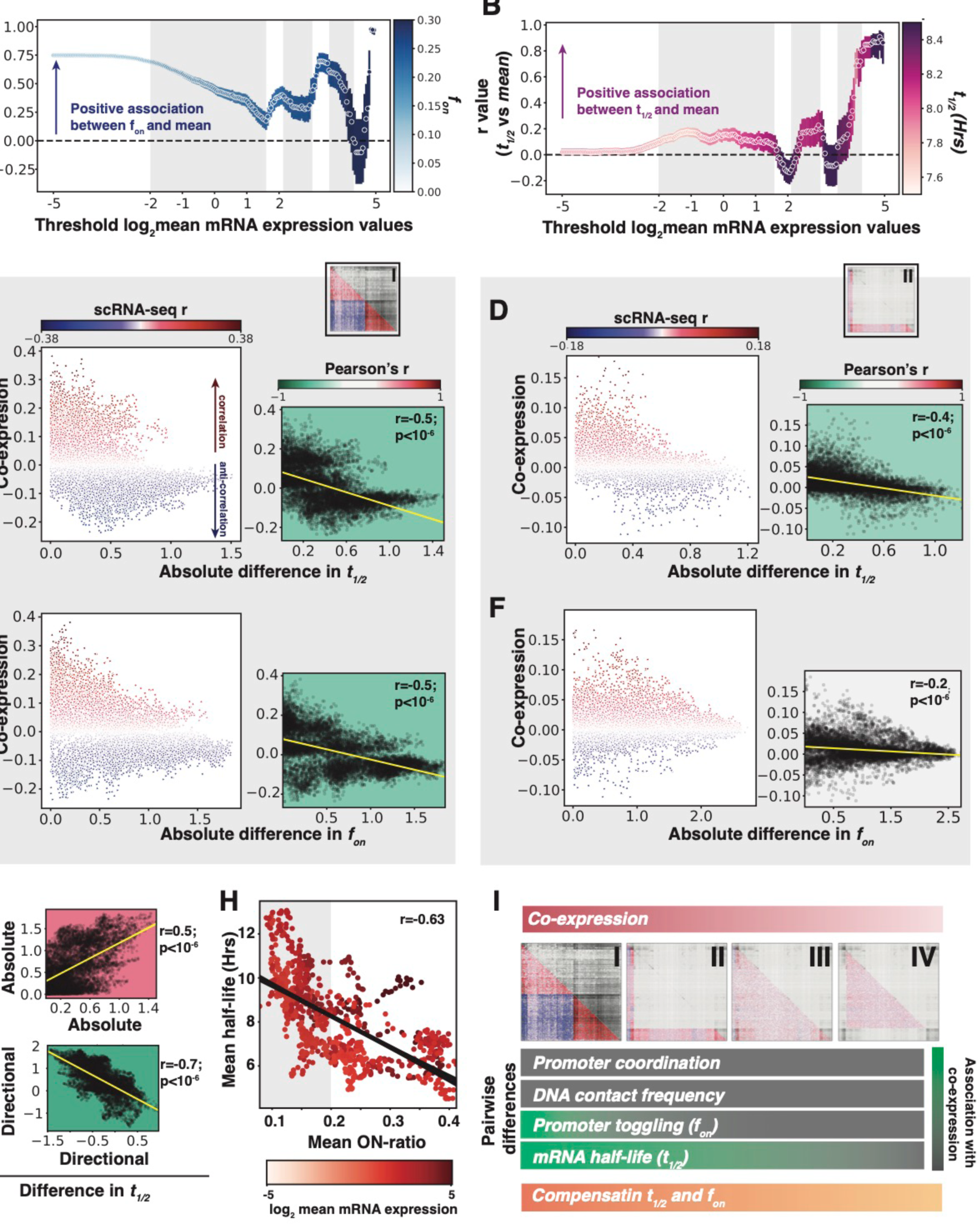
Differences in mRNA half-lives are most strongly associated with mRNA co-expression. **(A)** Quantification (Pearson’s r coefficient) of association between promoter ON ratio (*f*_*on*_) and relative mean mRNA expression (μ_log2_) calculated by imposing specific expression mean thresholds. Color bar represents the average promoter ON ratio (*f*_*on*_) for genes in each corresponding bin (error bar is 95% confidence interval). **(B)**Quantification (Pearson’s r coefficient) of association between transcript half-life (t_1/2_) and mean mRNA expression (μ_log2_) calculated by imposing specific expression mean thresholds. Color bar represents the average transcript half-life (t_1/2_) for genes in each corresponding bin (error bar is 95% confidence interval). **(C-D)** Scatter plot (left) and correlation (right) between average mRNA co-expression (i.e., scRNAseq Pearson’s r coefficient) and average difference in transcript half-life (C) or promoter ON ratio (D) for regime I. **(E-F)** Scatter plot (left) and correlation (right) between average mRNA co-expression (i.e., scRNAseq Pearson’s r coefficient) and average difference in transcript half-life (E) or promoter ON ratio (F) for regime II. **(G)**Top: correlation between average absolute difference in transcript half-life and average absolute difference in promoter ON ratio. Bottom: correlation between average directional difference in transcript half-life and average directional difference in promoter ON ratio. Sampling region corresponds to regime I. **(H)**Scatter plot showing correlation between average ON ration and half-lives for genes where log_2_(μ)>1. Dots are colored according to log_2_(μ) to compare with Figure S2H. **(I)**Schematic illustrating that sampled region with decreasing average mRNA co-expression behavior (i.e., regimes I to IV, left to right) show a decreasing correlation between mRNA co-expression and mRNA half-life.

To identify if pairwise differences in promoter toggling kinetics (*f*_*on*_) or mRNA half-lives (*t1/2*) underlie co-expression of genes at an mRNA level, we followed the same workflow as in Figure 1 and Figure S1. We performed random subsampling of regimes I-IV (Figure 1C), and plotted the relationship between pairwise mRNA co-expression (i.e., scRNA-seq r) and either the pairwise difference in mRNA half-life (Figure 2C and 2D) or the pairwise difference in the ON-fraction of the respective promoters (Figure 2E and 2F). In the regime with the highest mRNA co-expression (i.e., most positive and most negative scRNA-seq r values, regime I), both the promoter toggling and the mRNA half-life differences have a strong relationship with mRNA co-expression (Figure 2C and E). Nevertheless, this relationship becomes less prominent for promoter toggling when we sample regimes with more subtle mRNA co-expression at an mRNA level (Figure 2F, S2E and S2F). Conversely, pairwise differences in mRNA half-lives remain more strongly associated with mRNA co-expression even at lower co-expression levels (Figure 2D, S2C, and S2D). Together, these data show that mRNA degradation is coupled to both positive and negative co-expression across all ∼5000 genes analyzed, where similar half-lives are associated with positive co-expression, while different half-lives are associated with negative co-expression of two genes (Figure 2I).

### mRNA half-life and promoter toggling demonstrate compensatory behavior

The finding that both pairwise differences in promoter toggling as well as mRNA half-life are associated with mRNA co-expression for a small subset (<10%) of genes (Figure 2C and E), and seem to have an inverse effect on mean mRNA abundance (Figure 2A and 2B), led us to question the relationship between promoter toggling and mRNA half-life. Therefore, we quantified both the absolute and directional pairwise differences in half-life and promoter toggling (Figure 2G and S2G, see STAR Methods for details). In the regime with the highest co-expression values (regime I), there is a strong positive correlation between absolute differences in promoter toggling and mRNA half-life (Figure 2G and S2G, top). This means that gene-pairs with a large difference in promoter toggling kinetics, also display a large difference in mRNA half-life. Across all sampled regimes (I-IV) half-life and promoter toggling showed compensatory (i.e., genes with higher *f*_*on*_ have lower *t*_*1/2*_) rather than synergistic (i.e., the groups of genes with a higher *f*_*on*_ also demonstrate a higher *t*_*1/2*_) behavior (Figure 2G and S2G, bottom). Intuitively, this compensation can be explained by the inherent relationship between promoter toggling and mRNA half-life (*μ* = *f*_*on*_.*k*_*tx*_*/k*_*d*_, where *k*_*d*_ is mRNA degradation rate). Therefore, at similar mean mRNA expression levels we expect mRNA half-life to be inversely proportional to promoter toggling (Figure S2H). This relationship between two seemingly distal kinetic parameters emphasizes the balance between transcriptional and post-transcriptional processes that together orchestrate the gene expression landscapes, to the extent that the one cannot be understood without the other.

## DISCUSSION

In this brief report we integrated gene co-expression data from scRNA-seq analysis with other data modalities (Nora et al., 2017; Shah et al., 2018; Sharova et al., 2009) that provide context from various processes involved in gene expression to assess whether specific kinetic steps influence scRNA-seq data. Our goal was to interrogate the relationships between the observed correlations in single-cell mRNA levels and the processes upstream and downstream of transcription without assuming specific regulatory architectures. The analysis showed that promoter coordination was not playing a discernible role in gene co-expression especially when (anti)correlation of mRNA is the strongest (Figure 1). Yet, it is important to note that promoter behavior might not be properly captured by a single snapshot, as promoter coordination could emerge from more complex temporal dynamics where promoters are, for example, likely to be ON in close temporal proximity but not necessarily at the same time. Unfortunately, the techniques for assessing promoter activity with temporal resolution (e.g. MS2 tagging) are limited to lower throughput applications (Hocine et al., 2013; Wan et al., 2021) and require genetic modification, as a specific sequence has to be added to a gene for the mRNA to be tracked (Tantale et al., 2016).

We next included other kinetic parameters in the analysis. Strikingly, mRNA half-life demonstrated a strong negative association with scRNA-seq co-expression r values (Figure 2). Additionally, for specific regimes of very evident (anti)correlation in mRNA levels, differences in promoter ON ratio also seemed to play an important role. In fact, in these regions the differences in *f*_*on*_ and half-life demonstrate clear compensatory behavior. It is possible that there is a mechanistic reason why this occurs --i.e., is it a requirement for genes to have extreme correlation values to show this specific behavior --or that this relationship is an evolutionary consequence. Many elegant studies on gene expression regulation focus on chromatin and transcriptional events (Larson et al., 2013; Lenstra et al., 2016; Zinani et al., 2022), and downstream processes should not be underestimated. This work together with other recent publications (Aizarani et al., 2019; Gilbertson et al., 2018; Hansen et al., 2018; Hansen and Weinberger, 2019; Matkovic et al., 2022) is enforcing the necessity of including post-transcriptional events.

## ACKNOWLEDGEMENTS

We thank all members of the Hansen lab for the thoughtful discussion and suggestions. We also thank Hendrik Marks for his generous gift of mESC-E14 cell line. M.M.K.H. acknowledges support from the Dutch Research Council (NWO) ENW-XS awards (OCENW.XS3.055 and OCENW.XS21.2.050).

## AUTHOR CONTRIBUTIONS

Ó.G.B. and M.M.K.H. conceived the study. Ó.G.B., P.V., B.M, and M.M.K.H. designed the study. Ó.G.B. performed single-cell RNA-seq experiments. Ó.G.B., P.V., and M.M.K.H. designed and performed the data analyses. Ó.G.B. and M.M.K.H. wrote and P.V. and B.M. edited the manuscript

## DATA AND MATERIALS AVAILABILITY

Sequencing data from single-cell RNA-seq will be deposited onto GEO. Custom code for analysis of sequencing data and mathematical modeling will be made available on GitHub.

## STAR METHODS

### Key resources table

| REAGENT or RESOURCE | SOURCE | IDENTIFIER |
| --- | --- | --- |
| **Cell lines** |  |  |
| mESC E14 type (129/Ola background) | (van Mierlo et al., 2019) |  |
| **Cell culture reagents / utensils** |  |  |
| 6-well Clear TC-treated Multiple Well Plates | Costar | REF 3516 |
| Gelatin | Sigma-Aldrich | 48723/500g |
| Dulbecco's Modified Eagle Media, high glucose, pyruvate (Gibco) | Fisher Scientific | #11594486 |
| Beta-mercaptoethanol | Fisher Scientific | #11528926 |
| Sodium pyruvate (Gibco) | Fisher Scientific | #11530396 |
| Penicillin/Streptomycin (Gibco) | Fisher Scientific | #15140122 |
| ESGRO recombinant Mouse Leukemia Inhibitory Factor | Merckmillipore | ESG1107 |
| Fetal Bovine Serum | Fisher Scientific | #A3840002 |
| Phosphate-buffered saline (PBS) (Gibco) | Fisher Scientific | #10728775 |
| 0.05% Trypsin-EDTA (Gibco) | Fisher Scientific | #11500636 |
| DMSO | Sigma-Aldrich | D5879-1L |
| Acridine Orange/Propidium Iodide Stain | Logos Biosystems | F23001 |
| LUNA Cell Counting Slides | Logos Biosystems | L12001 |
| LUNA-FL Dual Fluorescence Cell Counter | Logos Biosystems | L20001-LG |
| **Deposited data** |  |  |
| Intron-seq FISH mESC | (Shah et al., 2018) |  |
| Hi-C contact frequencies mESC | (Nora et al., 2017) |  |
| Half-life mRNA mESC | (Harova et al., 2009) |  |
| scRNA-seq mESC | Produced in house |  |
| **Software and algorithms** |  |  |
| Seurat V3 | (Stuart et al., 2019) |  |

## RESOURCE AVAILABILITY

Further information and requests for resources and reagents should be directed to and will be fulfilled by the lead contact, M.M.K.H.

## METHODS DETAILS

### Single-cell RNA sequencing in mESCs

mESC-E14 (mESCs) in this study were obtained from Hendrik Marks’s group at Radboud University. mESC were seeded per well in gelatin coated Costar® 6-well Clear TC-treated Multiple Well Plates (REF 3516), in serum/LIF culture media consisting of Dulbecco’s Modified Eagle Media, supplemented with 0.1mM beta-mercaptoethanol, 1000 U/mL of Penicillin, 0.1 mg/mL of Streptomycin, 1mM of sodium pyruvate, 1000 U/mL of ESGRO recombinant Mouse Leukemia Inhibitory Factor and 15% of ES-qualified heat inactivated Fetal Bovine Serum. Cells were grown for 30 hours in a 37°C incubator with 5% CO_2_ atmosphere. Cells were exposed to 0.007% EtOH for the last 6 hours. Cells were then washed once with PBS and detached with 0.05% trypsin-EDTA for 4 minutes at 37°C. Detached cells were pelleted and resuspended in 1mL of freezing media consisting of 80% culture media, 10% of extra heat inactivated FBS and 10% of DMSO. Proper viability of the cells prior to freezing was quantified with propidium iodide/acridine orange staining in a LUNAs-FL Dual Fluorescence Cell Counter. Frozen cells were delivered to the commercial company Single Cell Discoveries (https://www.scdiscoveries.com) in dry ice. Single cell barcoding was performed in a 10x microfluidic genomic chip in order to encapsulate individual cells in water droplets in oil, containing cell specific barcoded beads. Labeled RNA molecules were pooled and subjected to a poly-A specific reverse transcription. cDNA molecules where linearly amplified using an *in vitro* transcription reaction and the final sequencing library was obtained through a second step of reverse transcription. 3’-end sequencing was performed, followed by quality control and genomic mapping. Final read count per gene and per cell matrices where generated and used in posterior analysis steps.

### Calculation of gene-to-gene correlation in scRNAseq data and gene clustering

Preprocessing of raw counts was performed with SeuratV3 (Stuart et al., 2019). Genes detected in less than 5 cells and cells with less than 200 detected genes, less than 24000 total RNA count and cells with more than 5% mitochondrial RNAs were discarded. Cells were given a score reflecting their cell cycle stage based on the expression of specific cell cycle markers (Kowalczyk et al., 2015). A normalized and scaled count matrix was generated by using the SCTransform method (Hafemeister and Satija, 2019) with an offset of 8=100 (Lause et al., 2021). The scaling method accounted for the total RNA count per cell and utilized the percent of mitochondrial RNA and the cell cycle scoring as variables to regress. The scaled count matrix was used to calculate the full correlation matrix using the Pearson or the Spearman methods. Pairwise gene-to-gene Pearson correlation matrices were clustered using complete-linkage hierarchical agglomerative clustering.

### Gene-to-gene promoter coordination calculation

From the intron-seq FISH data (Shah et al., 2018), the status of the promoter was assessed according to the presence or absence of intronic signal. Promoter ON states are defined by the presence of intronic signals (i.e., signal ≥ 1), while OFF states are defined by the lack thereof (i.e., signal = 0). Therefore, per gene:

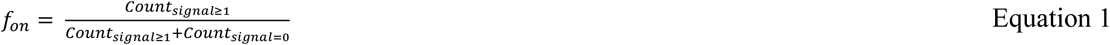

For each pair of genes in the dataset, an observed anti-coordination score (O) was defined as the proportion of cell pairs where the genes show an opposite behavior (ON/OFF or OFF/ON) over the total number of cell-to-cell comparisons as described in equation 1. The observed anti-coordination scored was corrected by the chance of these behavior appearing from independently behaving promoters. To this extent, the proportion of cells in which each gene is ON was defined as its *f*_*on*_, and the expected anti-coordination score (E) was calculated as described in equation 2, where 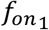 and 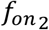 represent the ON ratios (*f*_*on*_) of each of the two genes included in the calculation. Observed anti-correlation score was corrected by calculating the percentual fold change of the observed anti-coordination score with respect to the expected score by chance as described in equation 3, where *O* is the observed anti-coordination score and *E* is the expected anti-coordination score. In order to present this data in a more intuitive way, we transformed the corrected anti-coordination score in a coordination score as described in equation 5.

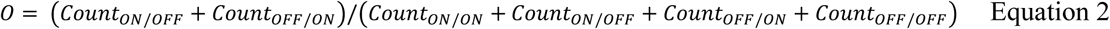

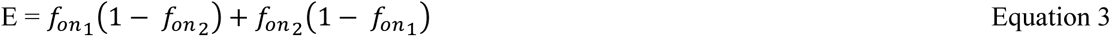

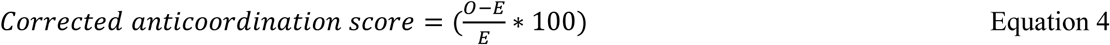

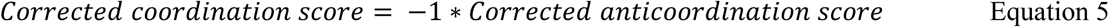

### Estimation of the relationship between promoter toggling rates

The effective ratio between *K*_*on*_ and *K*_*off*_ is calculated from the *f*_*on*_ quantification (Equation 1) where:

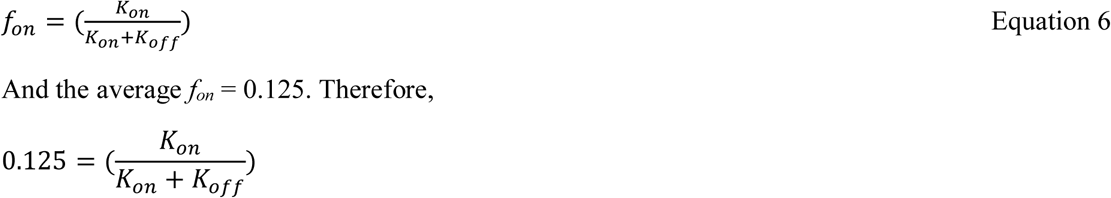

And the average *f*_*on*_ = 0.125. Therefore,

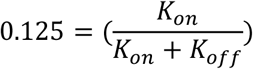

and,

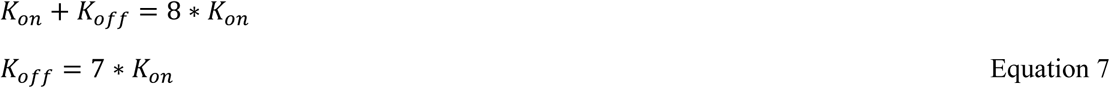

### Calculation of gene expression mean and noise from scRNA-seq dataset

Single gene mean and noise values were obtained through the analysis of the raw read count matrix with SeuratV3 (Stuart et al., 2019). Next, genes detected in less than 5 cells and cells with less than 200 detected genes, less than 24000 total RNA count and cells with more than 5% mitochondrial RNAs were discarded. Cells were given a score reflecting their cell cycle stage based on the expression of specific cell cycle markers (Kowalczyk et al., 2015). Mean gene expression values and residual variance values (*σ*^*2 res*^) were calculated using the SCTransform method with an offset value of 8=100, using the percent of mitochondrial RNA and the cell cycle score as variables to regress. In short, biological noise is obtained through the fitting of the read counts to a negative binomial distribution modeling technical noise and obtaining the residual variance not explained by the model of technical noise (Hafemeister and Satija, 2019). These residual variance values (o^2 *res*^) are comparable to the Fano factor (Lause et al., 2021). Mean gene expression was then calculated as the log^2^(μ) denoted as *μ*^*log2*^.

### Rolling averages and analysis of correlation for gene expression parameters

#### Subsampling

For the four sampling regimes, we randomly sub-sampled 10000 sets of 30 by 30 genes (sampling regimes II, III and IV) or 3 by 3 genes (sampling regime I), within each defined region. Sampling regime I is a much smaller region so we scaled down the subsampling accordingly.

#### Rolling averages

Genes were sorted according to the matrix resulting from the clustering of the pairwise scRNA-seq Pearson’s coefficients. Starting from the first ranked gene, groups of successive 30 genes were made in an iterative fashion moving 1 position forward in the gene list with each iteration. For each group of 30 genes the average of *fon, μ*^*log2*^, *σ*^*2*^_*std*_ and *t*_*1/2*_ was obtained. This process generated four sequential vectors of averages in which each position corresponded to the same group of genes from the scRNA-seq clustering order.

#### Correlation for gene expression parameters

To analyze the correlation between the kinetic parameters (*fon* and *t*_*1/2*_) and the gene expression output parameters (μ_log2_ and σ^2^_std_), Pearson coefficients as well as the 95% confidence intervals for the Pearson coefficients were calculated between the corresponding pairwise values for each parameter in the previously calculated rolling mean vectors. In order to classify gene groups in different regimes, they were grouped in bins according to their μ_log2_ average value by using a rolling threshold for this value.

### Binned analysis of correlations between differences in kinetic parameters

Two random groups of 30 or 3 contiguous genes in the clustered scRNAseq r matrix were selected in order to form a 30×30 or 3×3 comparative space. Restrictions in the selection of gene groups were applied in order to create the 4 sampling regions described previously. For each selected space, the average r coefficient was calculated. By matrix symmetry the average promoter coordinated behavior for the corresponding group of genes was obtained from the processed intron-seqFISH data and the average Hi-C contact frequency was obtained with the same method from the Hi-C data. The average difference in mean, noise, ON ratio and half-life (i.e., *μ*_*log2*_, *σ*^*2*^_*std*_, *fon*, and *t*_*1/2*_) was obtained by averaging these parameters for each of the two groups in the comparison and obtaining the directional or absolute difference as described in equations 8 and 9 respectively.

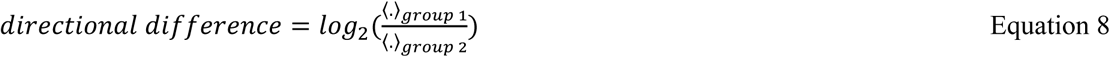

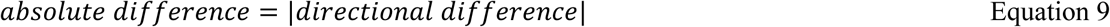

## FIGURE LEGENDS

**Figure S1:**
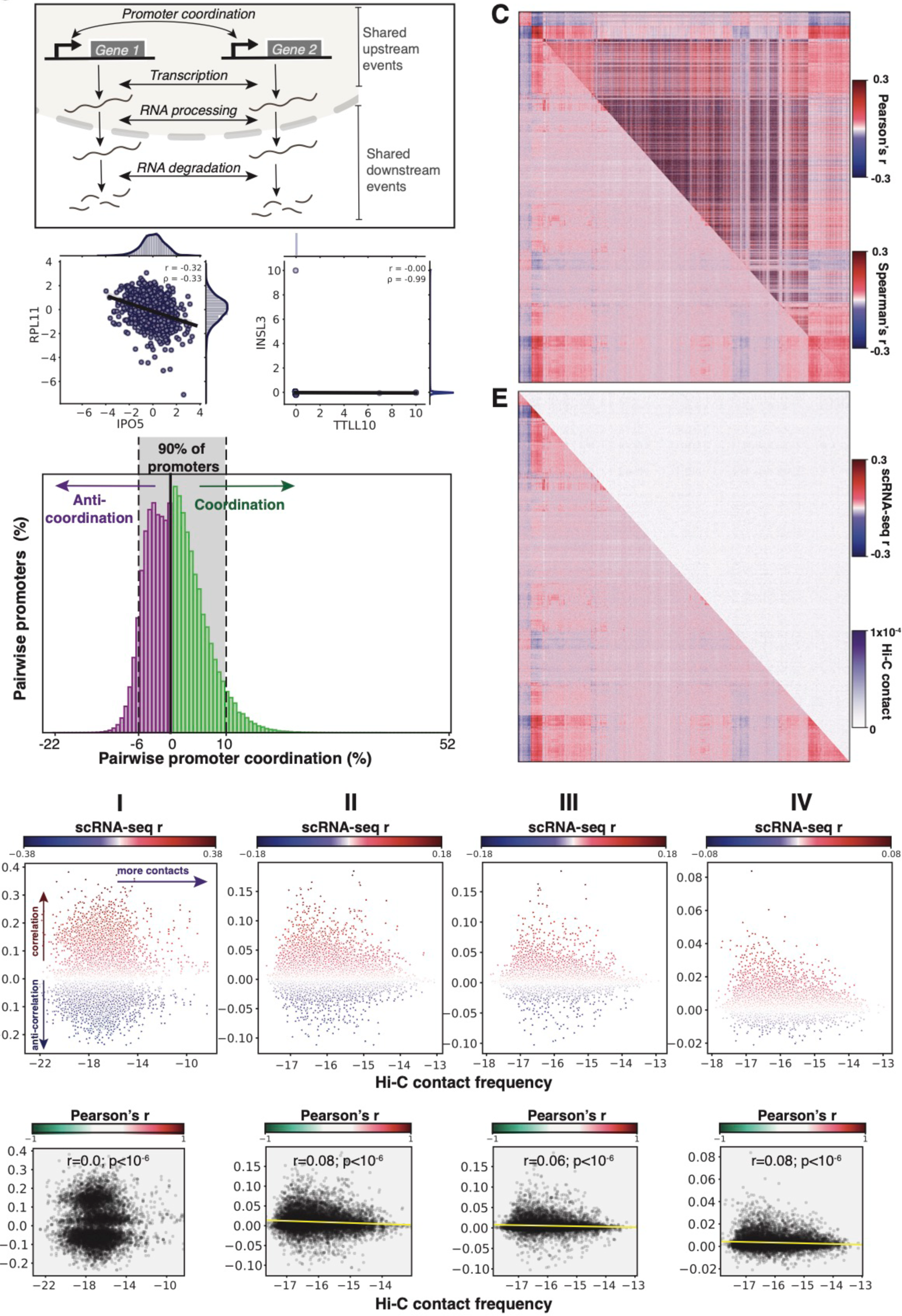
Gene-to-gene contacts do not explain mRNA co-expression (Related to Figure 1). **(A)**Schematic illustration of two genes that can exhibit mRNA co-expression by a shared transcriptional (upstream) kinetic step or a shared post-transcriptional (downstream) kinetic step. **(B)**Scatter plots representing scaled single cell mRNA expression values for pairs of genes where both Pearson and Spearman correlation coefficients give the same r values (left), and where Spearman coefficients are unreliable (right). **(C)**Combined matrix showing clustered gene-to-gene scRNAseq expression Pearson r coefficients (bottom half) and corresponding Spearman coefficients (top half). **(D)**Frequency of pairwise promoter interactions that are either more coordinated (green) or more anti-coordinated (purple) than expected from random promoter behavior. Shaded area represents 90% of pairwise comparisons. **(E)**Combined matrix showing clustered gene-to-gene scRNAseq expression Pearson’s r coefficients (bottom half) and corresponding Hi-C contact frequencies (top half). **(F-G)** mRNA co-expression (i.e., scRNAseq Pearson’s r coefficient) versus average Hi-C contact frequencies for regions I, II, III and IV. **(F)** Color (blue to red) represent the average scRNAseq Pearson r coefficient. **(G)** Correlation between mRNA co-expression and Hi-C contact frequencies, with linear regression line plotted in yellow and the background color corresponding to Pearson’s r correlation coefficient.

**Figure S2:**
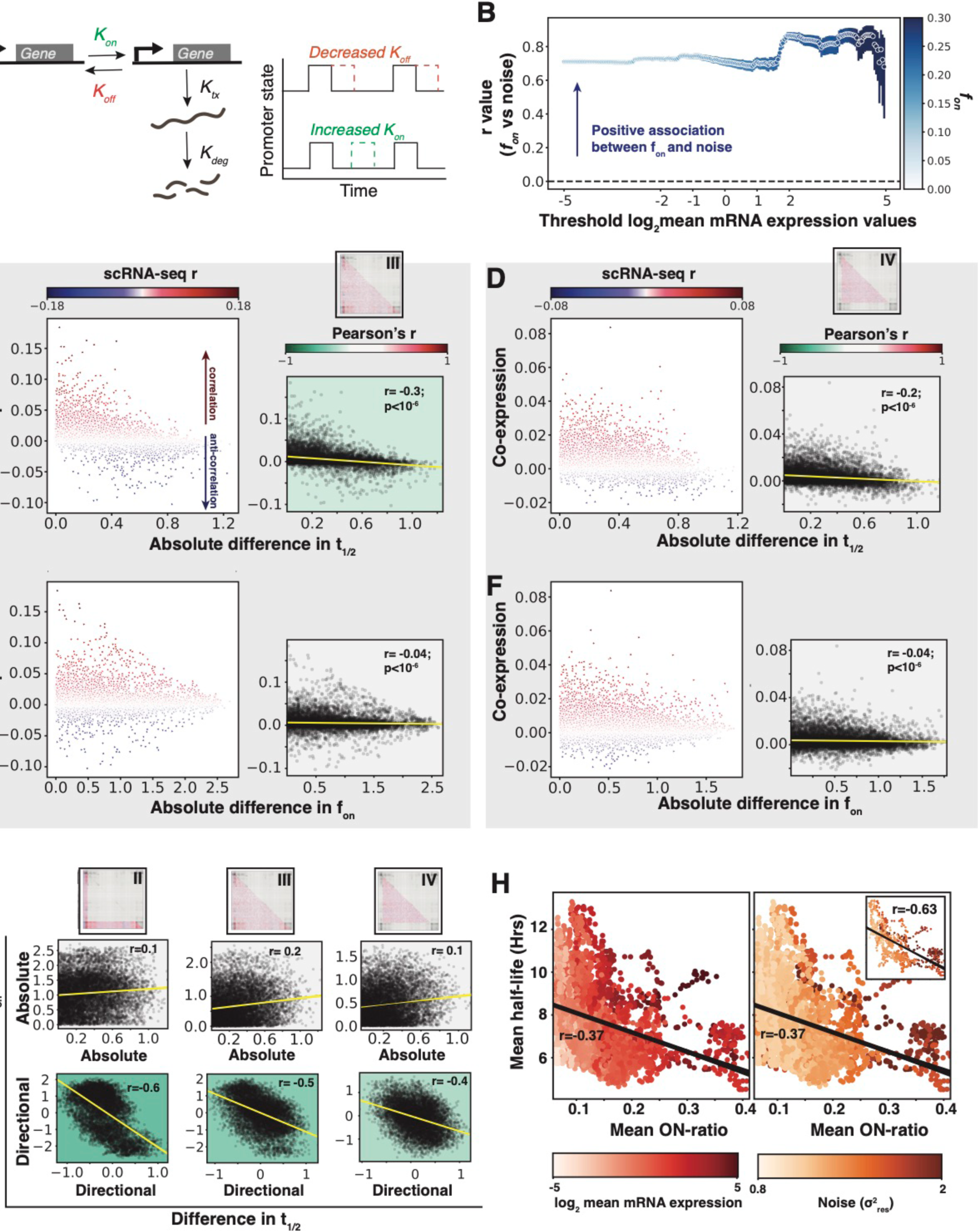
Differences in promoter toggling does not associate with mRNA co-expression and shows compensation behavior with mRNA degradation (Related to Figure 2). **(A)**Schematic illustrating how promoter toggling can be impacted by altering either the switching ON of a promoter (*K*_*on*_) or the switching OFF of a promoter (*K*_*off*_). **(B)**Quantification (Pearson’s r coefficient) of association between promoter toggling (*f*_*on*_) and relative mRNA noise (Fano, μ_log2_) calculated by imposing specific expression mean thresholds. Color bar represents the average promoter toggling (*f*_*on*_) for genes in each corresponding bin (error bar is 95% confidence interval). **(C-D)** Scatter plot (left) and correlation (right) between average mRNA co-expression (i.e., scRNAseq Pearson’s r coefficient) and average difference in transcript half-life (C) or promoter ON ratio (D) for regime III. **(E-F)** Scatter plot (left) and correlation (right) between average mRNA co-expression (i.e., scRNAseq Pearson’s r coefficient) and average difference in transcript half-life (E) or promoter ON ratio (F) for regime IV. **(G)**Top: correlation between average absolute difference in transcript half-life and average absolute difference in promoter ON ratio. Bottom: correlation between average directional difference in transcript half-life and average directional difference in promoter ON ratio. Sampling region corresponds to regime II-IV from left to right. **(H)**Scatter plot showing correlation between average ON ration and half-lives per gene. Dots colored according to log_2_(μ) (left) or noise (right). Inset: Scatter plot showing correlation between average ON ration and half-lives for genes where log_2_(μ)>1. Dots colored according to noise.

